# Female-biased embryonic death from genomic instability-induced inflammation

**DOI:** 10.1101/400820

**Authors:** Adrian J. McNairn, Chen-Hua Chuang, Jordana C. Bloom, Marsha D. Wallace, John C. Schimenti

## Abstract

Genomic instability (GIN) can trigger cellular responses including checkpoint activation, senescence, and inflammation. Though extensively studied in cell culture and cancer paradigms, little is known about the impact of GIN during embryonic development, a period of rapid cellular proliferation. We report that GIN-causing mutations in the MCM2-7 DNA replicative helicase render female mouse embryos to be dramatically more susceptible than males to embryonic lethality. This bias was not attributable to X-inactivation defects, differential replication licensing, or X vs Y chromosome size, but rather “maleness,” since XX embryos could be rescued by transgene-mediated sex reversal or testosterone (T) administration. The ability of exogenous or endogenous T to protect embryos was related to its anti-inflammatory properties. The NSAID ibuprofen rescued female embryos containing mutations not only in MCM genes but also *Fancm,* which have elevated GIN from compromised replication fork repair. Additionally, deficiency for the anti-inflammatory IL10 receptor was synthetically lethal with the GIN-causing *Mcm4*^*Chaos3*^ helicase mutant. Our experiments indicate that embryonic and maternal GIN arising from DNA replication-associated DNA damage induces embryonic inflammation likely via the cGAS-STING response, preferentially killing female embryos while male embryos are protected by high levels of intrinsic T.

---

Compromised DNA replication or replication-associated repair can lead to GIN. Such replicative stress (RS) can be induced environmentally or by intrinsic genetic defects such as oncogene expression or mutation of DNA replication and repair genes ^1,2^. Consequences of RS can trigger the DNA damage response (DDR), causing a delay or halt in cell cycle progression that is particularly impactful for rapidly proliferating cells ^3^. Embryogenesis constitutes one of the most dramatic examples of rapid and choreographed cell proliferation, thus GIN can potentially have profound effects upon a fetus. Cells experiencing chronic RS, GIN, or DNA damage can exhibit a “senescence-associated secretory phenotype” (SASP), typified by secretion of several proteins including inflammatory cytokines that can inhibit growth of neighboring cells in a culture system ^4^. SASP is related to the cGAS/STING pathway that induces an inflammatory response triggered by the presence of cytoplasmic DNA ^5^. Though senescence and the SASP have normal functions in certain compartments during embryonic development ^6,7^, little is known about potential consequences of fetal or maternal GIN-induced inflammation during gestation.

DNA replication requires the minichromosome maintenance (MCM) complex, composed of 6 proteins (MCM2-7) that constitute the catalytic core of the replicative helicase. Experimental reduction of MCMs causes RS by reducing the number of dormant (“backup”) origins that are important for completing DNA replication in the face of spontaneous or induced replication fork stalling or collapse ^8–10^. We previously described a hypomorphic allele of the mouse gene *Mcm4* (*Mcm4*^*Chaos3*^; abbreviated *Mcm4*^*C3*^ hereafter) that was identified in a screen for mutants with elevated micronuclei (membrane-encapsulated cytoplasmic chromosome fragments) ^11^. Highly cancer-prone, this allele causes GIN by destabilizing the MCM2-7 helicase and triggering TRP53-mediated reduction (∼40%) of MCM2-7 in mouse embryonic fibroblasts (MEFs) ^12–14^. Although *Mcm4*^*C3*^ homozygotes in the C3H strain background are fully viable, further reduction of MCM levels, achieved *via* heterozygosity for certain other *Mcm* genes, caused severe phenotypic consequences including pre-and postnatal lethality and growth defects ^12^. Upon closer examination of those and additional breeding data, we noticed that females of the MCM-depleted, semi-lethal genotypes *Mcm4*^*C3/Gt*^, *Mcm4*^*C3/C3*^ *Mcm2*^*Gt/+*^,*Mcm4*^*C3/C3*^*,Mcm6*^*Gt/+*^, and *Mcm4*^*C3/C3*^ *Mcm7*^*Gt/+*^ (Gt = gene trap allele) were drastically under-represented (up to several fold) compared to males of the same mutant genotype (Fig. 1a; Tables S1-S4). There was no gender skewing associated with non-lethal genotypes (*Mcm4*^*C3/C3*^ *Mcm3*^*Gt/+*^ and *Mcm4*^*C3/C3*^ *Mcm5*^*Gt/+*^), i.e., those genotypes present in offspring at Mendelian ratios (Fig. 1a; Table S5) ^12^.

**Fig. 1.**
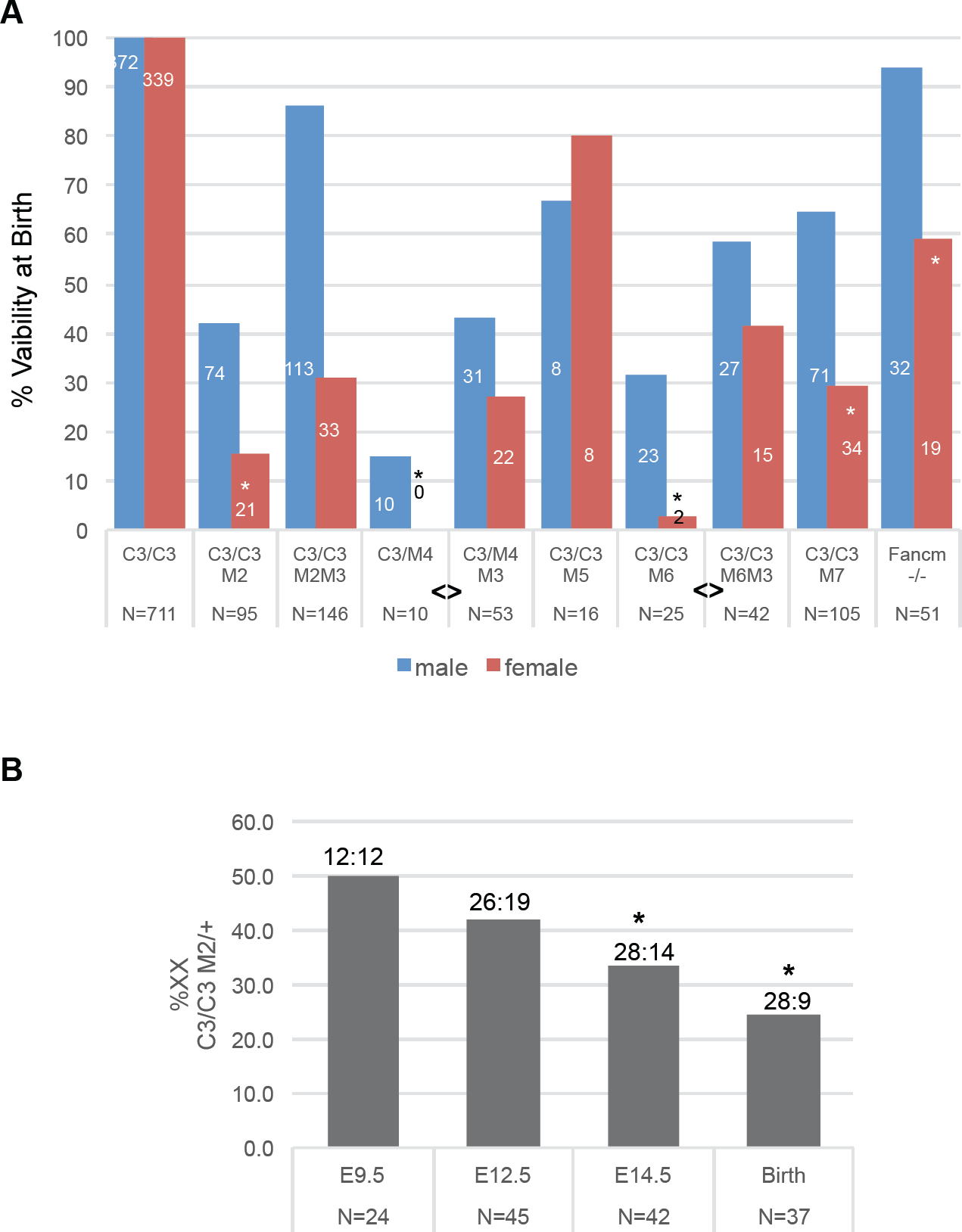
Female-biased embryonic lethality in MCM-depleted mice. **(A)** Female MCM-depleted mice are underrepresented at birth. Mice were bred to produce *Mcm4*^*C3/C3*^ (“C3/C3”) or *Mcm4*^*Chaos3/Gt*^ offspring (“C3/M4), some of which were heterozygous for null alleles in other MCMs (“M#”; for example, M2/+ = *Mcm2*^*Gt/+*^). Graphed are percent viability at birth of males and females for each of the indicated genotypes versus C3/C3 littermates. For *Fancm*, the viability is versus WT littermates. The numbers on or over the bars = # males or females of -the indicated genotype, and the N values below equal the total number of newborns with that genotype. Some of the data for all genotypes except that involving M5 were reported in ^12^, but broken out by sex here and with added data that are enumerated in Extended Data Table 1 and Tables S1-S5. “*” represents a Chi-Squared test probability of p<0.01 that females are underrepresented vs males from that genotype. “<>” represents a significant Fisher Exact Test (p<0.05) between indicated groups in terms of the ability of *Mcm3* heterozygosity to decrease sex bias. **(B)** Timing of female death during embryogenesis. E = embryonic day. Numbers above bars are viable XY:XX embryos genotyped (they sum to the “N” values). Asterisks indicates a Chi-Squared probability of p<0.05.

To determine the stage(s) of development that *Mcm4*^*C3/C3*^ *Mcm2*^*Gt/+*^ animals were dying and if this coincided with female-biased lethality, we conducted timed matings of *Mcm4*^*C3/C3*^ females to *Mcm4*^*C3/+*^ *Mcm2*^*Gt/+*^ males, and genotyped embryos at different points of gestation and at birth. Loss of *Mcm4*^*C3/C3*^ *Mcm2*^*Gt/+*^ embryos (compared to *Mcm4*^*C3/C3*^) was first evident at E14.5, and was already skewed against females; the male:female ratios of *Mcm4*^*C3/C3*^ *Mcm2*^*Gt/+*^ embryos at E9.5, E12.5, E14.5 and birth were 1.0, 1.37, 2.0, and 3.11, respectively (Fig. 1b).

Previous studies showed that in MEFs derived from *Mcm* mutant mice, decreased MCM levels correlated with a reduction in dormant replication origins ^15,16^. Activation of licensed dormant origins near stalled/collapsed replication forks is important for mitigating RS ^9^. To test if decreased dormant origin licensing contributes to female-biased lethality in our models, *Mcm3* heterozygosity was introduced into the embryonic lethal genotypes. Hypothesized to have a role in exporting nucleoplasmic MCMs into the cytoplasm, *Mcm3* heterozygosity was shown to ameliorate *in vivo* and *in vitro* phenotypes (viability through gestation; cancer susceptibility; growth) of MCM-deficient mutant mice and cells by increasing chromatin-bound MCMs ^12^. The introduction of *Mcm3* heterozygosity dramatically rescued viability of *Mcm4*^*C3/Gt*^ and *Mcm4*^*C3/C3*^ *Mcm6*^*Gt/+*^ female embryos preferentially, increasing female viability from 0% to 27% in the former, and from 3% to 42% in the latter (Fig. 1a, Tables S1,S2). *Mcm3* heterozygosity also increased viability of *Mcm4*^*C3/C3*^ *Mcm2*^*Gt/+*^ newborns from 30% to 72%, but both sexes were rescued approximately proportionately (Fig. 1a; Table S3).

We conjecture that the degree of preferential female rescue is related to overall degree of lethality in compound mutants (93%, 82% and 70% lethality for *Mcm4*^*C3/Gt*^, *Mcm4*^*C3/C3*^ *Mcm6*^*Gt/+*^ and *Mcm4*^*C3/C3*^ *Mcm2*^*Gt/+*^, respectively) ^12^.

We hypothesized that the biased lethality of female embryos under conditions of *Mcm* deficiency was related to one of the following: 1) defects in X-inactivation, 2) the larger size of the X (∼171 Mb) vs the Y chromosome (∼90 Mb), or 3) secondary sexual characteristics. Regarding #1, it was conceivable that impaired or delayed DNA replication might disrupt inactivation of one X chromosome, leading to severe developmental defects ^17^. To test this, we bred *Mcm4*^*C3/+*^ *Mcm2*^*Gt/+*^ males bearing an ubiquitously-expressed X-linked GFP transgene ^18^ to *Mcm4*^*C3/C3*^ females. Mid-gestation (E10.5) female embryos, all of which must bear the X-linked GFP transgene, were analyzed by flow cytometry to determine the fraction of GFP+ cells. There was no difference between non-lethal genotypes (*Mcm4*^*C3/+*^ *Mcm2*^*Gt/+*^*; Mcm4*^*C3/+*^ *Mcm2*^*+/+*^*; Mcm4*^*C3/C3*^ *Mcm2*^*+/+*^) and the sex-biased lethal genotype (*Mcm4*^*C3/C3*^ *Mcm2*^*Gt/+*^; Extended Data Fig. 1), indicating that X-inactivation occurs normally.

Next, to distinguish between hypotheses 2 and 3 (whether the larger size of the X *vs* secondary sexual characteristics might underlie the sex bias), a single experiment was performed. Sex determination occurs from E9.5-12.5, when the bipotential gonad differentiates down the male or female pathway ^19^. During this interval, expression of the Y-linked sex determining gene *Sry* is critical for initiating testis development by triggering the differentiation of precursor cells into pre-Sertoli cells ^20^. We induced sex reversal of XX *Mcm4*^*C3/C3*^ *Mcm2*^*Gt/+*^ embryos with an autosomal *Sry* transgene (*Sry129 Tg4Ei*) ^21^. Strikingly, presence of the *Sry* transgene increased the proportion of XX *Mcm4*^*C3/C3*^ *Mcm2*^*Gt/+*^ mice from 20% to 48% (Fig 2a; Table S6). These results indicate that maleness, and not the presence of two X chromosomes *per se*, protects embryos from MCM deficiency. These data are also consistent with the finding that preferential female embryo death only occurred after sex determination.

**Fig. 2.**
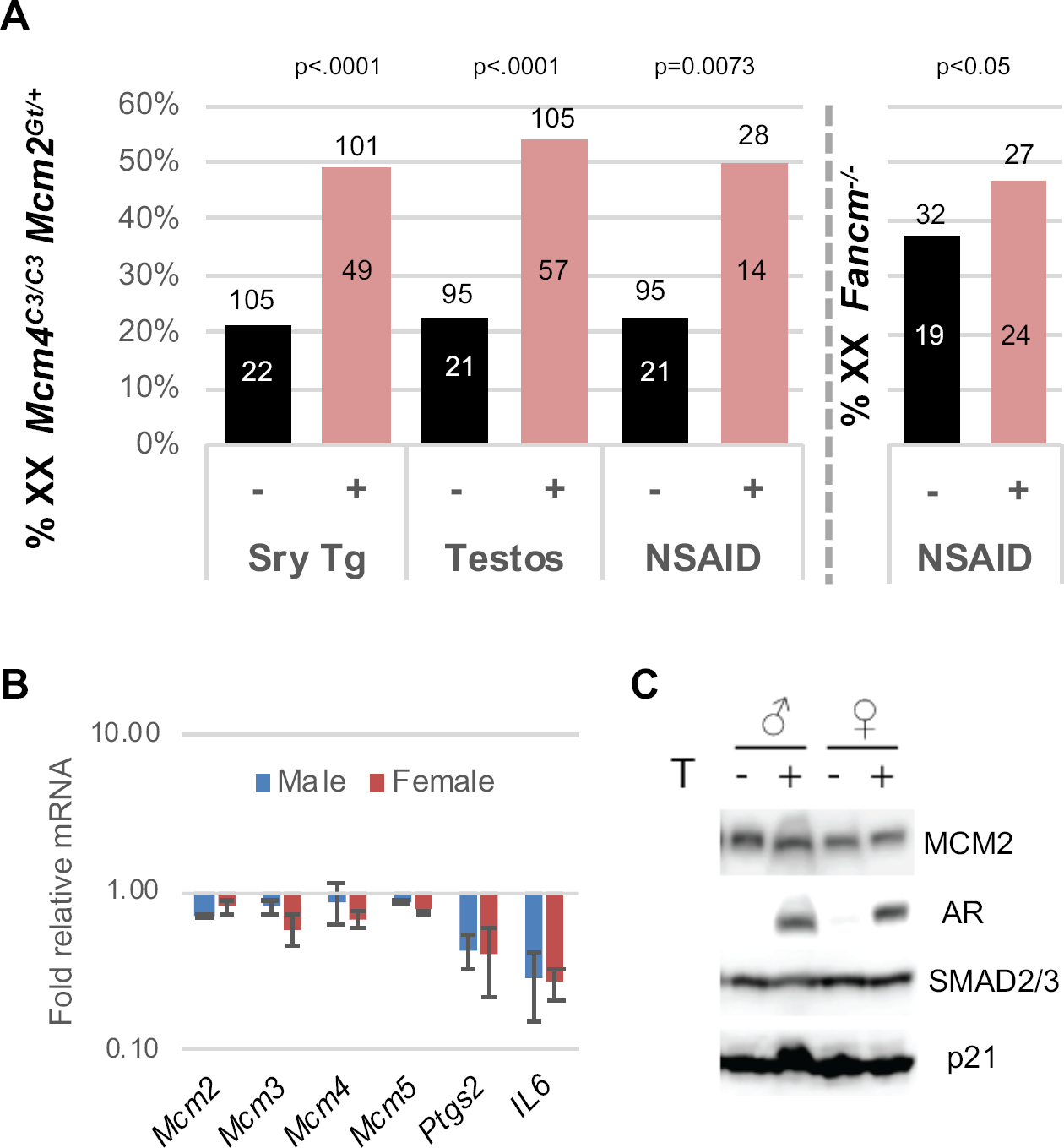
Evidence that the anti-inflammatory activity of testosterone protects male embryos from genomic instability-induced lethality. **(A)** Viability of genetically female (XX) *Mcm4*^*C3/C3*^ *Mcm2*^*Gt/+*^ or *Fancm*^*-/-*^ embryos is rescued by *Sry* transgene-induced sex reversal (Sry Tg), and treatment of pregnant dams with either testosterone (“Testos”) or ibuprofen (“NSAID”). Values above each bar are total mice, and those inside bars are XX. Nontransgenics and transgenics in the Sry Tg experiment were from the same cross. The untreated mice in the testosterone and NSAID experiments are from Table S3 and also plotted in Fig. 1a. This aggregate value contains 8 male and 1 female *Mcm4*^*C3/C3*^ *Mcm2*^*Gt/+*^ offspring that were produced contemporaneous to the NSAID cohort. P values are from two-tailed F.E.T. The Sry Tg and testosterone crosses involved an *Mcm3*^*Gt*^ allele that was included to boost viability (but not sex skewing, see Fig. 1a) of the lethal genotypes. **(B)** Testosterone treatment does not affect *Mcm* mRNA, but does lower the inflammation markers *Il6* and *Ptgs2*. Following T treatment (10nM) of *Mcm4*^*C3/C3*^ *Mcm2*^*Gt/+*^ MEFs were T treated for 1 hr, and mRNA collected 6 hrs later for qRT-PCR. **(C)** Same as B, but protein was collected 24 hrs after T treatment, and Western analysis performed with indicated antibodies.

Since sex reversal rescued lethality of XX embryos, we hypothesized that testosterone (T) might be responsible. It is produced at high levels by Leydig cells in embryonic testes, beginning at ∼E12.5 and persisting throughout gestation ^22^. To test this, we treated pregnant females bearing embryos of the *Mcm4*^*C3/C3*^ *Mcm2*^*Gt/+*^ genotype with daily injections of testosterone propionate beginning at E7.5. Fetuses were collected and genotyped at E19.5 (T-treated dams cannibalized newborns). The viability of XX animals from T-treated mothers (XX females were masculinized by T) increased dramatically compared to the untreated embryos (Fig. 2a; Table S7), rising from 22% to 54% for this genotype.

Regarding the mechanism by which T protects untreated male and T-treated female embryos from GIN, we first considered the possibility that it does so by increasing replication licensing. It was reported that the androgen receptor, to which T and its more potent metabolite dihydrotestosterone binds, stimulates proliferation of prostate cancer cells by acting as a replication factor ^23,24^. However, there were no sex-specific differences in MCM2 or MCM4 protein levels in E13.5 fetuses or placentae of various genotypes (Extended Data Fig. 2). Additionally, we observed no increase of *Mcm* mRNA or chromatin-bound MCMs in T-treated mouse embryonic fibroblasts (MEFs) (Fig. 2b, c). Next, we hypothesized that T was functioning by ameliorating certain consequences of the elevated GIN in the *Mcm* mutants. In particular, chromosome damage-induced formation of micronuclei, a signature phenotype of *Mcm4*^*Chaos3*^ mice ^11^, can trigger inflammation via the cGAS-STING pathway ^25^. T, a steroid hormone, suppresses the expression of pro-inflammatory cytokines while increasing the anti-inflammatory molecule IL10 (interleukin 10) ^26–29^. Indeed, T treatment of *Mcm4*^*C3/C3*^ *Mcm2*^*Gt/+*^ MEFs caused >2-3 fold decreases in *Il6* (interleukin 6, a pro-inflammatory cytokine) and *Ptgs2* (the human ortholog of *COX2*, encoding cyclooxygenase 2, that is central for production of prostaglandins that cause inflammation and pain) mRNAs (Fig. 2b). Next, to more directly test if inflammation impacts the sex bias in *Mcm4*^*C3/C3*^ *Mcm2*^*Gt/+*^ mice, we treated pregnant females beginning at 7.5 days of gestation with ibuprofen in their drinking water, then genotyped their newborns. Strikingly, ibuprofen completely abolished the sex bias, decreasing the male (XY) to female (XX) sex ratio from 2:1 to 1:1 without altering secondary sex characteristics (Fig. 2a, Extended Data Table 1a). The NSAID treatment did not affect MCM protein levels in embryos or placentae (Extended Data Fig. 2b, c).

The findings that two distinct anti-inflammatory molecules (one steroidal, the other not) rescued XX embryos is consistent with the interpretation that GIN and resulting elevated micronuclei caused by the DNA replication defects cause inflammation that leads to preferential female embryonic lethality. Nevertheless, because ibuprofen and the androgen receptor (which is strongly induced by T exposure; Fig. 2c) can alter expression of genes unrelated to inflammation ^30–32^, we used an orthogonal approach to further test the idea that inflammation underlies embryonic lethality in the MCM mutants. We hypothesized that genetically increasing inflammation by ablating the receptor (*Il10rb*) for the anti-inflammatory cytokine IL10 would exacerbate the phenotype of *Mcm4*^*C3/C3*^ *Mcm2*^*+/-*^ lethality. In the course of breeding such mice, we found that the genotype of *Mcm4*^*C3/C3*^ *Il10rb*^*-/-*^ caused highly penetrant lethality to embryos of both sexes (Extended Data Table 1c). IL10 mediates a feedback loop under conditions of inflammation to induce degradation of *Ptgs2*/*Cox2* transcripts ^33^, and also counters the inflammation response triggered by the STING pathway ^34^. To confirm that the synthetic lethality of *Il10r*^*-/-*^ with *Mcm4*^*C3/C3*^ was due to inflammation, we treated pregnant females with NSAID during gestation. This rescued both males and females of the *Mcm4*^*C3/C3*^*Il10rb*^*-/-*^ genotype, increasing viability from 8.9% to 94% (Extended Data Table 1c). These results indicate that the level of GIN in certain *Mcm4*^*C3/C3*^ tissues, even in the absence of addition MCM deletions, can cause inflammatory responses that can be lethal unless attenuated by endogenous or exogenous anti-inflammatories.

Successful pregnancy requires suppression of inflammation at the maternal:fetal interface. Because homozygosity for *Chaos3* alone causes a ∼20 fold increase in micronucleated erythrocytes without decreasing viability in the C3H background^11^, and IL10 is thought to play a role in suppressing maternal inflammation at the fetal:maternal interface ^35^, we considered the possibility that maternal genotype might influence viability of MCM compound mutant embryos. To test this, we mated females heterozygous for *Chaos3* (*Mcm4*^*C3/+*^ *Mcm2*^*Gt/+*^) to *Mcm4*^*C3/C3*^ males (all data presented heretofore were from *Mcm4*^*C3/C3*^ females X *Mcm4*^*C3/+*^ *Mcm2*^*Gt/+*^ males) and genotyped the offspring. Surprisingly, this cross abolished the sex bias against *Mcm4*^*C3/C3*^ *Mcm2*^*Gt/+*^ females at birth, indicating that the maternal genotype impacts the viability of *Mcm-* deficient embryos (Fig. 3a; Extended Data Table 1b).

**Fig 3.**
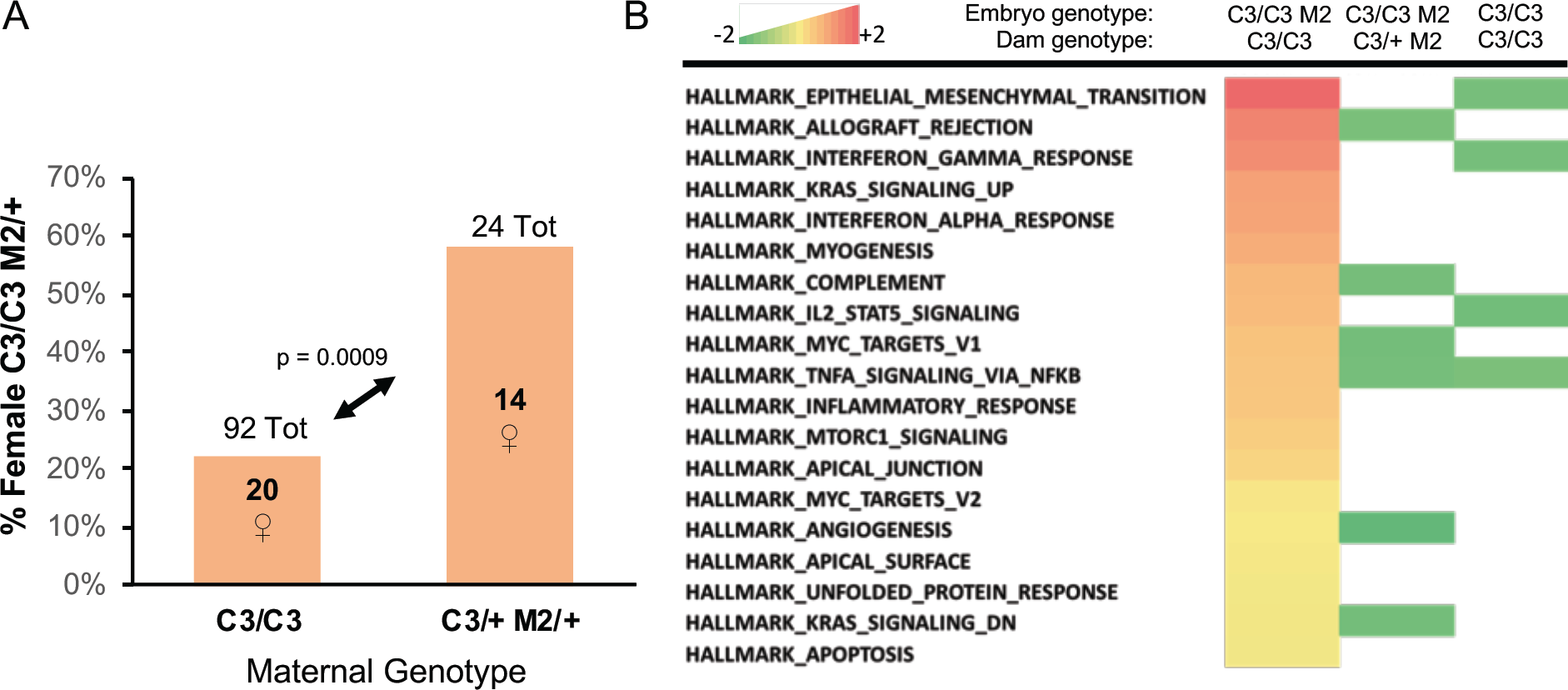
Maternal GIN genotype impacts female embryo viability and placental inflammation. **(A)** Lethality of female (XX) *Mcm4*^*C3/C3*^ *Mcm2*^*Gt/+*^ embryos is dependent upon maternal genotype. C3 = *Mcm4*^*C3*^; M2 = *Mcm2*^*Gt/+*^. The p-value was calculated by Fisher’s Exact Test. See Tables S2 and Extended Data Table 1b for primary data from the crosses. **(B)** Placentae of female E13.5 embryos with the *Mcm4*^*C3/C3*^ *Mcm2*^*Gt/+*^ lethal genotype have elevated expression of inflammation pathways when the dam has elevated GIN. Shown are heatmaps of GSEA analysis of RNA-seq data, using the Hallmarks dataset of the Molecular Signatures Database (MSigDB; http://software.broadinstitute.org/gsea/msigdb/collections.jsp). Only those Hallmark pathways that were significantly different between sexes (FDR <0.25, nominal p value<0.05) were used to generate the heatmap. Multiple pathways involving inflammation are upregulated in *Mcm4*^*C3/C3*^ *Mcm2*^*Gt/+*^ female vs male embryos from *Mcm4*^*C3/C3*^ dams, but not other combinations. Embryonic and maternal genotypes are listed at the top of the heatmaps.

These data led us to hypothesize that maternal homozygosity for *Chaos3* imposes additional stress on the placentae of genetically susceptible female embryos, possibly via the cGAS/STING-driven inflammation. To test this, we examined DNA damage levels in placentae of E13.5 embryos produced in control (WT x WT) and reciprocal *Mcm4*^*C3/+*^ *Mcm2*^*Gt/+*^ X *Mcm4*^*C3/C3*^ and *Mcm4*^*C3/C3*^ X WT crosses. Regardless of fetal genotype, placentae from embryos in which the dam was *Mcm4*^*C3/C3*^ had more cells that stained positively for the double strand break (DSB) marker γH2AX than those in which the dam was of any other genotype (Fig. 4a). NSAID treatment did not reduce the level of γH2AX staining, consistent with the rescue effect being related to inflammation, not DNA damage or GIN *per se* (Fig. 4a).

**Fig. 4.**
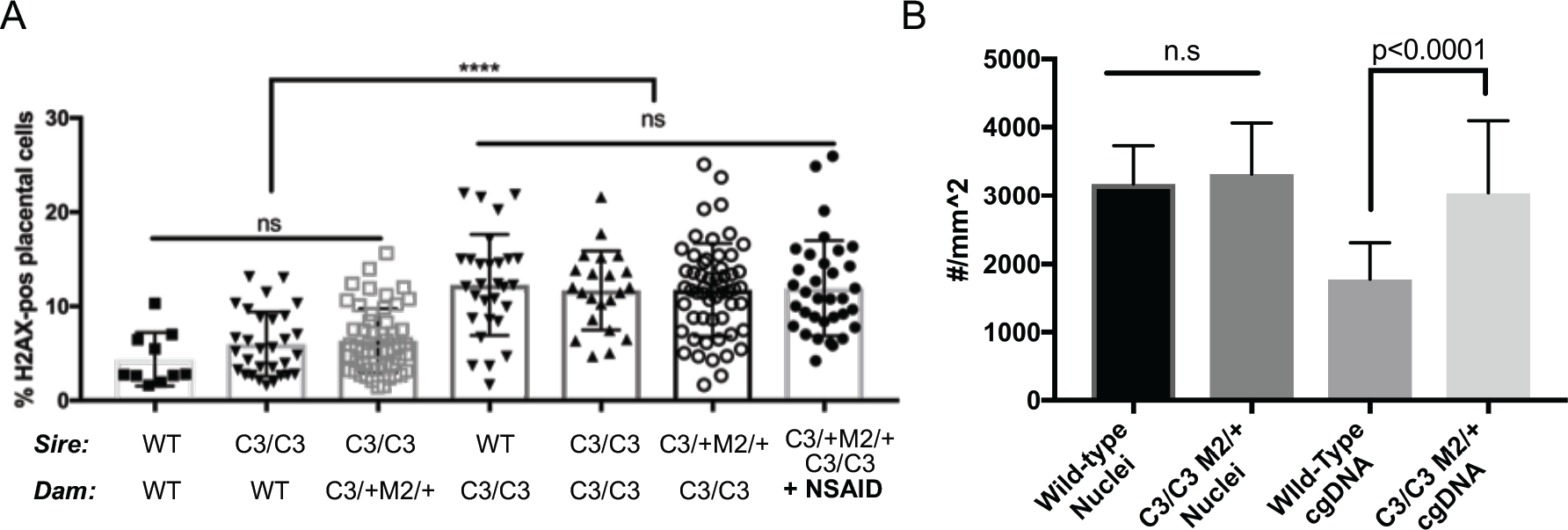
Dams with intrinsic GIN cause elevated DNA damage in the placenta. **(A)** γH2AX staining in placentae from the indicated maternal genotypes. Each dot represents a single placenta. Significance was by unpaired, 2-tail t-tests. Error bars = standard deviation. ns = not significant. **(B)** *Mcm4*^*C3/C3*^ *Mcm2*^*Gt/+*^ placentae have elevated cytoplasmic DNA. Cytoplasmic DNA was determined by staining E13.5 placentas with an anti-dsDNA antibody and quantifying multiple sections of each placenta (10 *Mcm4*^*C3/C3*^ *Mcm2*^*Gt/+*^ and 10 wild-type). The numbers on the graph represent total number of nuclei and DNA foci (“cgDNA”) per mm^2^ of a slide section. The p-value is from a two-tailed t-test between WT and mutant. C3 = *Mcm4*^*C3*^; M2 = *Mcm2*^*Gt/+*^; WT = strain C3H +/+.

Given the maternally-induced DNA damage in the placenta and the evidence that inflammation is responsible for the preferential death of females in our mouse model, we hypothesized that GIN-induced placental inflammation might underlie the lethality. We tested this in three ways. First, if this model is true, then it implicates the placenta as potentially being the culpable defective tissue responsible for embryonic death. To explore this, we examined MCM levels in the placentae vs. embryo proper of various genotype of embryos. We observed significant reductions in placental, but not embryonic MCM2 and MCM4 (especially MCM4) in *Chaos3* mutant genotypes, regardless of maternal genotype or whether the dams were NSAID-treated (Extended Data Fig. 2a-c). Thus, placental cells may be particularly sensitive to DNA replication defects that trigger downregulation of MCM production and consequent increases in GIN ^13,36^. Second, because the cGAS-STING pathway has emerged as the primary means by which GIN and cytoplasmic DNA (e.g. micronuclei) induces an innate immune response, we stained WT and *Mcm4*^*C3/C3*^ *Mcm2*^*+/-*^ E13.5 placentae with an anti-dsDNA antibody to quantify cytoplasmic DNA foci/micronuclei. Consistent with the decreased levels of MCMs, there were nearly 2 fold more cytoplasmic DNA foci in placentae of the lethal genotype, regardless of maternal genotype (Fig. 4b; Extended Data Fig. 3). Third, we performed RNA-seq analysis of male vs female placentae from either *Mcm4*^*C3/+*^*Mcm2*^*Gt/+*^ or *Mcm4*^*C3/C3*^ dams. Gene set enrichment analysis (GSEA) showed increased expression of hallmark inflammation gene sets only in the lethal genotype combination of *Mcm4*^*C3/C3*^ *Mcm2*^*Gt/+*^ females from *Mcm4*^*C3/C3*^ dams (Fig. 3b; Extended Data Fig. 4). The major upregulated hallmarks included EMT transition (which is commonly associated with inflammatory responses ^37^), allograft rejection, and interferon gamma response. All three of these categories contain genes involved in inflammation and the innate immune response (Extended Data Fig. 4). These results are consistent with the genetic data indicating that maternal GIN synergizes with females of replication-impaired genotypes to cause lethal levels of inflammation.

While the data presented thus far demonstrate that MCM depletion (e.g. *Mcm2*, *Mcm6* or *Mcm7* hemizygosity) in conjunction with the destabilized replicative helicase in *Chaos3* mice trigger inflammation and embryonic death, it is unclear exactly what defects are primarily responsible, and whether the sex-biased phenomena are entirely unique to these models. We therefore attempted to parse the key proximal defects that trigger the sex bias by exposing WT embryos to either exogenous RS alone or DSBs alone. Pregnant females treated with hydroxyurea, an agent that causes RS by depleting nucleotide pools, delivered pups without sex skewing (M:F 1.08, 127 live births, p-value=0.5396 using one-tailed Fisher Exact Test). Similarly, chronic exposure to ionizing radiation (50 mGy administered 3X/week beginning at E1.5) of pregnant females also failed to produce a sex bias (M:F 1.00, 32 live births). We then conjectured that cytoplasmic DNA arising specifically from chronic DNA replication defects might be responsible for inflammation-driven female-biased embryonic lethality. It was reported that mice with mutations in *Fancm* (Fanconi Anemia M), encoding a protein important for stabilization and repair of DNA replication forks, cause elevated erythrocyte micronuclei ^38^ and under-representation of homozygous female offspring in a mixed strain background ^39^. We generated a new allele of *Fancm* by CRISPR/Cas9 editing in the C57BL/6J embryos and, consistent with the published results, observed a reduction in *Fancm*^*-/-*^ females (M:F 1.68; χ^2^ p = 0.03; Fig 1a, Extended Data Table 1d). As with the *Mcm4*^*C3/C3*^ *Mcm2*^*+/-*^ model, ibuprofen treatment of pregnant females rescued the sex ratio (*Fancm*^*-/-*^ M:F 1.13 vs *Fancm*^*+/+*^ 1.07; Fig. 2a, Extended Data Table 1d). These results indicate that defective DNA replication and/or replication-associated DNA repair results in an inflammatory response (likely driven by elevated micronuclei) that preferentially kills female embryos.

In humans, the male:female birth rate is ∼1.05 ^40^. Studies of spontaneous abortuses from women with recurrent miscarriages have revealed a highly preferential loss of female fetuses unrelated to karyotypic abnormalities, ∼10 fold in one study ^41^ and 1.7 fold in another ^42^. The biological basis for the preferential sensitivity of post-sex-determination stage female embryos in humans is unknown. However, our results raise the possibility that ∼midgestation NSAID administration during pregnancies involving conditions of maternal inflammation may have protective effects for the fetus.

## Acknowledgements

We thank the Cornell Transgenic Core facility for help in producing the *Fancm* mutant mutant mice, and Jen Grenier from the RNA sequencing-seq Core. This work was supported by grants from NIH (R01 HD086609 and T32 HD057854 to JCS, the latter supporting MDW) and the Department of Defense (BC083376 to C-HC).

## Author Contributions

C-H. C. made the original observations of sex skewing and studies with T and the *Sry* transgene. A.M. discovered the link to inflammation, performed those related studies, and also the contribution of maternal genotype; M.W. performed the X inactivation study. J.B. developed the *FancM* mice and carried out those breedings. J.S. supervised all aspects of the work and wrote the manuscript.

## Methods

### Mouse husbandry

All breeding and husbandry all crosses were performed in the same animal facility and room at Cornell’s Veterinary College (East Campus Research Facility), and under the same environmental conditions and health status. Research was conducted with an IACUC-approved protocol to JCS (0038-2004).

### Testosterone Injections and Sex Reversal

*Mcm4*^*C3/C3*^ *Mcm2*^*Gt/+*^ males were mated to *Mcm4*^*C3/C3*^ females and 100uL of a 3mg/ml solution of testosterone propionate (Sigma) was injected sub-cutaneously into the hind leg of pregnant females daily from E7.5 to E.16.5 (20μg/g/day). This dose has been shown to increase female fetal testosterone by 80% in a rodent model without serious toxicological effect ^1^. The testosterone propionate was dissolved in corn oil and filter sterilized prior to injection. For treatment of MEFs, a 50mM solution of testosterone propionate was prepared in ethanol and cells were treated with 10nM for 1 hour. The media was then removed and the cells collected at indicated timepoints. Sex reversal of *XX Mcm4*^*C3/C3*^ *Mcm2*^*Gt/+*^ embryos was carried out using an autosomally-linked *Sry* transgene (Tg(*Sry129)4Ei*) ^2^.

### Ibuprofen Treatment

*Mcm4*^*C3/C3*^ *Mcm2*^*Gt/+*^ males were mated to *Mcm4*^*C3/C3*^ females and at E7.5-E9.5 the pregnant females were provided with water bottles containing ibuprofen (Children’s Advil) 5mL(100mg) in 250mL. They were allowed to drink ad libitum (50-80mg/kg/day). Newborns were genotyped at birth for sex with *Sry* primers and *Mcm* mutation status. Control mice utilized the same male with another *Mcm4*^*C3/C3*^ female and no drug treatment. For *Il10rb*, the strain was obtained from Jax Mice (stock#005027) and backcrossed into the C3Heb/FeJ(Jax stock#000658) background for six generations (N6) before crossing into the *Mcm4*^*C3*^ strain for 2 additional generations (N2). *Mcm4*^*C3/C3*^*Il10rb*^*+/-*^ males were mated to *Mcm4*^*C3/C3*^*Il10rb*^*+/-*^ females and provided with ibuprofen as described above. Newborns were genotyped with primers for *Il10rb* (Table S9).

### Genotyping

Genomic DNA was isolated from animal tissue using the hot-shot lysis procedure ^3^. Genotyping PCR was carried out using Taq and gene-specific primer pairs (Table S9). For *Chaos3* genotyping, the PCR products were digested with MboII to identify mutant alleles as *Chaos3* but not wild-type alleles are digestible with this enzyme. For *Mcm5*, ES cells were verified using primers containing regions outside of the gene trap insertion to verify. To determine the sex of early embryos, primers for *Sry* (Sex-determining region Y) were used to identify males, females are *Sry* negative. Genotyping for *Mcm2-7* genetraps has been previously described ^4^.

### Generation of *Mcm5* mutant mice

*Mcm5*^*tm1a(KOMP)Mbp*^ genetrap ES cells (Mcm5_F10, ESC#477873) were obtained from the Mouse Biology Program (MBP) at UC Davis and injected into B6(Cg)-*Tyr*^*c-2J*^*/J* blastocyst donors to generate chimeras. Disruption of *Mcm5* was confirmed by PCR as described in genotyping section (Table S9). Following germline transmission, the mutation was backcrossed into C3H for ≥ 4 generations before crossing to C3H-*Mcm4*^*Chaos3*^ mice.

### Generation of *FancM* mice

*Fancm*^*em1/Jcs*^ was generated using CRISPR/Cas9-mediated genome editing. In summary, an optimal guide sequence targeting the first exon of *Fancm* was designed using the mit.crispr.edu website. Oligos to generate the sgRNA DNA template were ordered from Integrated DNA Technologies (IDT) and the sgRNA was *in vitro* transcribed as described previously ^5^ (CRISPR-FancF: GAAATTAATACGACTCACTATAGGCCAGCTGGTAGTCGCGCACGGTTTTAGAGCTA GAAATAGC, CRISPR-FancR: CAAAATCTCGATCTTTATCGTTCAATTTTATTCCGATCAGGCAATAGT TGAACTTTTTCACCGTGGCTCAGCCACGAAAA). Embryo microinjection into C57BL/6J zygotes was performed as described previously ^6^ using 50ng/uL of sgRNA and 50ng/uL of Cas9 mRNA (TriLink Biotechnologies). The resulting 7bp deletion was identified via Sanger sequencing and subsequent genotyping was performed with primers sets specific to the mutant and wild-type alleles. (Table S9).

### Flow cytometry to monitor X-inactivation

A transgenic mouse containing an X-linked *EGFP* was obtained and crossed to *Mcm4*^*Chaos3*^ mice. FACS analysis of embryos was carried out as described ^7^. *Mcm4*^*C3/+*^ *Mcm2*^*Gt/+*^ males bearing an ubiquitously-expressed *X*-linked *GFP* transgene were bred to *Mcm4*^*C3/C3*^ females. E10.5 female embryos (littermates from 7 different pregnancies), all of which must bear the *GFP* transgene, were genotyped, dispersed into single cells, and analyzed by flow cytometry to determine the fraction of GFP+ cells. Theoretical maximum of GFP-positive cells in controls is 50%.

### Quantitative real-time reverse transcription-PCR (qRT-PCR)

RNA was isolated from cells using a kit per manufacturer’s instructions (Zymo or Qiagen RNeasy). 500ng of RNA was reverse transcribed into cDNA using qScript (Quanta) and analyzed on an ABI7300 or a Bio-Rad CFX96 using the following primers and iTaq (Bio-Rad). All reactions were normalized to *Gapdh* and/or *Tbp*. Primer sequences are available in Table S10. *Il6, Ptgs2, Mcm2, Mcm3, Mcm4, Mcm5*.

### RNA-seq

Total RNA was isolated from E13.5 placentas by homogenizing placentas in RNA lysis buffer followed by column purification per manufacturers’ instructions(Omega Biotech). RNA sample quality was confirmed by spectrophotometry (Nanodrop) to determine concentration and chemical purity (A260/230 and A260/280 ratios) and with a Fragment Analyzer (Advanced Analytical) to determine RNA integrity. Ribosomal RNA will be subtracted by hybridization from total RNA samples using the RiboZero Magnetic Gold H/M/R Kit (Illumina). Following cleanup by precipitation, rRNA-subtracted samples were quantified with a Qubit 2.0 (RNA HS kit; Thermo Fisher). TruSeq-barcoded RNAseq libraries were generated with the NEBNext Ultra II Directional RNA Library Prep Kit (New England Biolabs). Each library was be quantified with a Qubit 2.0 (dsDNA HS kit; Thermo Fisher) and the size distribution was be determined with a Fragment Analyzer (Advanced Analytical) prior to pooling. Libraries will be sequenced on a NextSeq500 instrument (Illumina). At least 20M single-end 75bp reads were generated per library. For analysis, reads were trimmed for low quality and adaptor sequences with cutadapt v1.8 using parameters: -m 50 -q 20 –a AGATCGGAAGAGCACACGTCTGAACTCCAG --match-read-wildcards. Reads were mapped to the mouse reference genome/transcriptome using tophat v2.1 with parameters: --library-type=fr-firststrand --no-novel-juncs -G <ref_genes.gtf>. For gene expression analysis: cufflinks v2.2 (cuffnorm/cuffdiff) was used to generate FPKM values and statistical analysis of differential gene expression ^8^. For the GSEA analysis, all expressed genes were analyzed using the Hallmarks dataset ^9^. The placental gene sets used were comparisons between male and female *Mcm4*^*C3/C3*^ *Mcm2*^*Gt/+*^ placentae from *Mcm4*^*C3/C3*^ dams or *Mcm4*^*C3/+*^ *Mcm2*^*Gt/+*^ dams, and male versus female *Mcm4*^*C3/C3*^ from *Mcm4*^*C3/C3*^ dams.

### Immunoblotting

Protein was isolated from E13.5 placentas and embryos by acetone precipitation from RNA-isolation buffer (Buffer RLT or TRK) and resuspending in SUTEB loading buffer (8M Urea, 1% SDS, 10mM EDTA, 10mM Tris-HCl, pH 6.8). Protein lysates were run on 4-20% SDS-PAGE acrylamide gels and transferred to PVDF membrane (Millipore). Immunoblots were probed with anti-Mcm2 (Epitomics/Abcam), anti-MCM2(Cell Signaling Technology), anti-androgen receptor (Epitomics/Abcam), anti-SMAD2/3(Cell Signaling), anti-p21(Santa Cruz), anti-MCM4(Cell Signaling Technology), anti-actin (Sigma). Secondary antibodies used included goat anti-rabbit-HRP (Cell Signaling) and goat anti-mouse-HRP (Sigma). Crescendo ECL substrate(Millipore) was used and immunoblots digitally scanned using a cDigit scanner. Quantification of immunoblots was performed using ImageStudio software.

### γH2ax and anti-dsDNA Staining

Placentae from E13.5 embryos were dissected from individual embryos and washed in PBS. Decidua were separated from placenta and uterine tissue with fine forceps. Genotyping was carried out using a piece of the embryo. Placentae were flash-frozen in OCT and 10μM sections cut on a cryostat and affixed to slides. Sections were fixed for 10 minutes with 4% paraformaldehyde in PBS, and stained with mouse anti-γH2ax-phospho ser41 (Millipore) or anti-dsDNA(Abcam) using a M.O.M kit and Biotin-Streptavidin blocking kit (Vector Labs) according to manufacturer’s instructions. Alexa-488 or Alexa 647-streptavidin (Invitrogen) was used to visualize. Slides were scanned using a Scanscope FL with a 20X objective. For cytoplasmic DNA detection, images were quantified using Fiji and foci were detected as described ^10^ with an added size parameter to differentiate between nuclei and cytoplasmic signals ^11^

### Hydroxyurea and IR treatment of embryos

For irradiation experiments, pregnant females were irradiated with 5 Rads (50mGy), 3 times a week during gestation, beginning at E1.5. For HU experiments, hydroxyurea (Sigma) was dissolved at 10mg/ml in sterile 1X PBS for injection. Pregnant C3H females were subjected to daily i.p. injections of 30-50ug/kg beginning at E3.5. Control females received daily i.p injections of sterile 1X PBS alone. All pregnancies were carried to term and the number and sex of animals determined at birth.

---

**Extended Data Figure 1.**
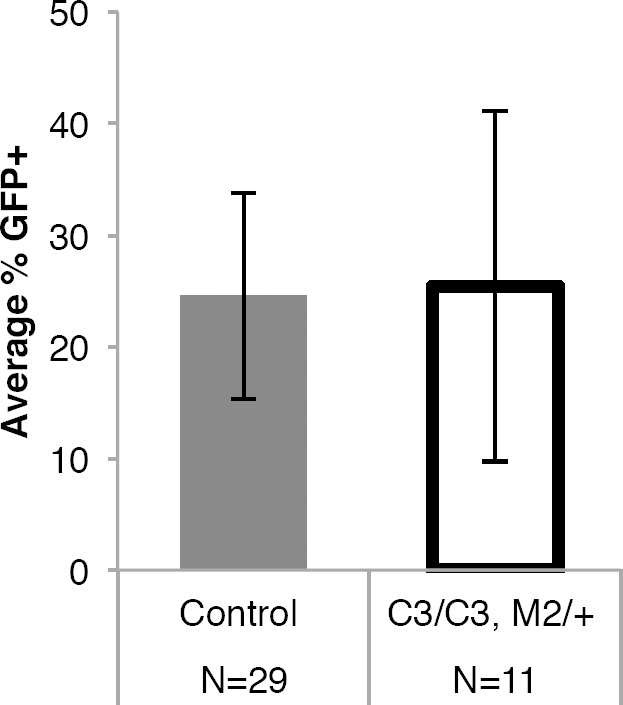
X-inactivation is not perturbed in MCM mutant embryos. Mouse female embryos bearing one X-linked GFP transgene were dispersed into single cells and examined by flow cytometry for GFP fluorescence. Control animals were female littermates with a genotype of *Mcm4*^*C3/+*^ *Mcm2*^*+/+*^ or *Mcm4*^*C3/C3*^ *Mcm2*^*+/+*^. The error bars represent the variability in GFP-positive cells among the individual embryos (N) used. There is no significance difference between the values by an unpaired T-test. C3 = *Mcm4*^*Chaos3*^; M2 = *Mcm2*^*Gt*^.

**Extended Data Figure 2.**
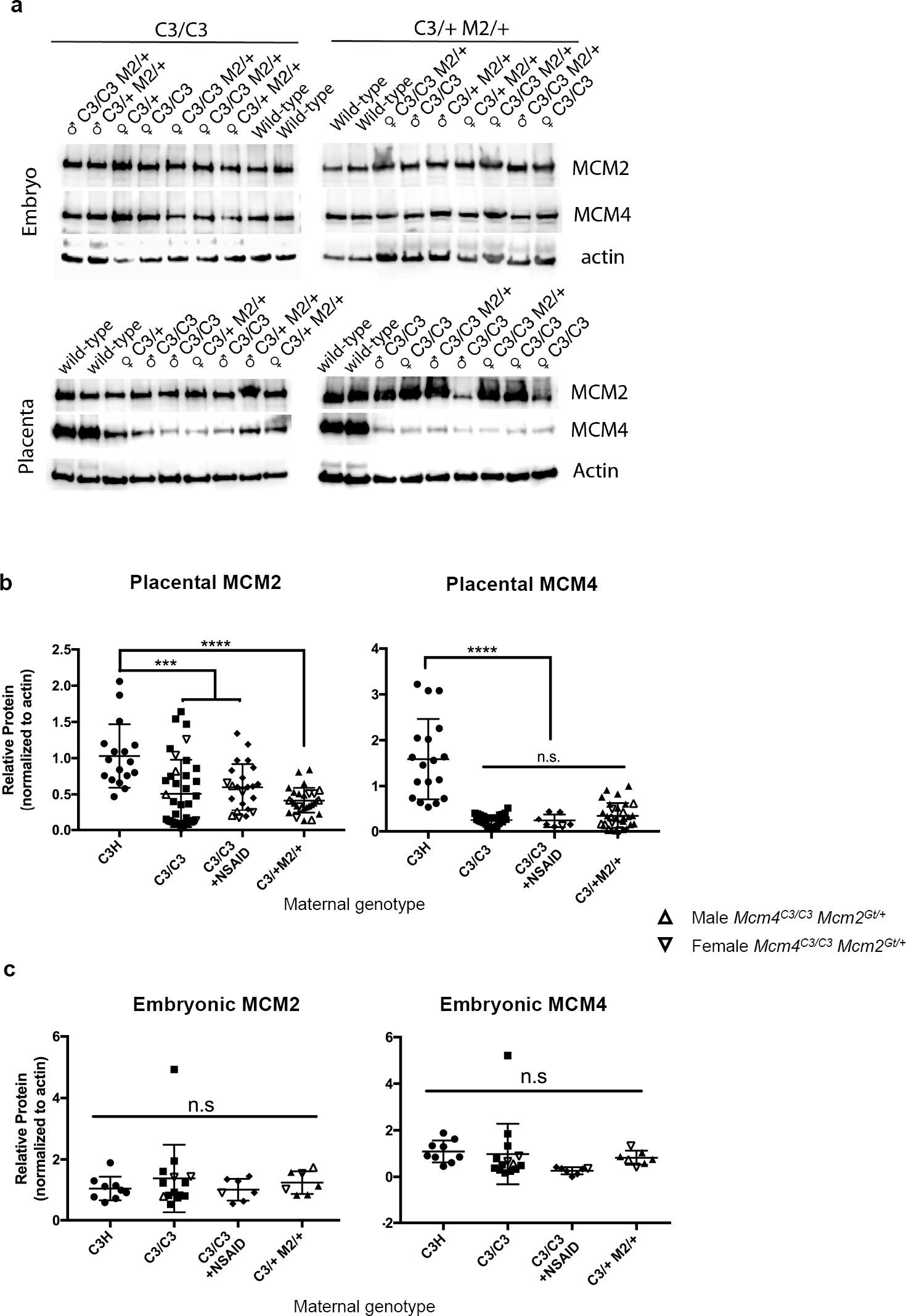
Placental, but not embryonic MCM levels are decreased in MCM mutants independent of maternal genotype, and NSAID does not rescue MCM levels. **a)** Representative westerns blots of protein lysates from E13.5 embryos and placentas of the indicated genotypes (top of each lane) were immunolabeled with antibodies against MCM2, MCM4, and beta actin. The samples came from dams of two genotypes indicated at the top of the panel. C3 = *Mcm4*^*Chaos3*^; M2 = *Mcm2*^*Gt*^. Note that MCM4 levels are particularly affected. **b)** Placental MCM2 and MCM4 protein levels from the indicated maternal genotypes were quantified from western blots (including some other than those in “a”) that were imaged (see Methods) and normalized to actin and WT protein levels. Each plotted point represents a single placenta. Asterisks indicate significance by unpaired two-tailed T-test (***, p<0.001,****, p<0.0001). Placentae corresponding to male or female *Mcm4*^*C3/C3*^ *Mcm2*^*Gt/+*^ genotype are indicated.**c)** Embryonic MCM2 and MCM4 protein levels were determined as in (**b**). Each plotted point represents a single embryo. Embryos corresponding to male or female *Mcm4*^*C3/C3*^ *Mcm2*^*Gt/+*^ genotype are indicated. The results were not significant (n.s.) by a one-way ANOVA.

**Extended Data Figure 3.**
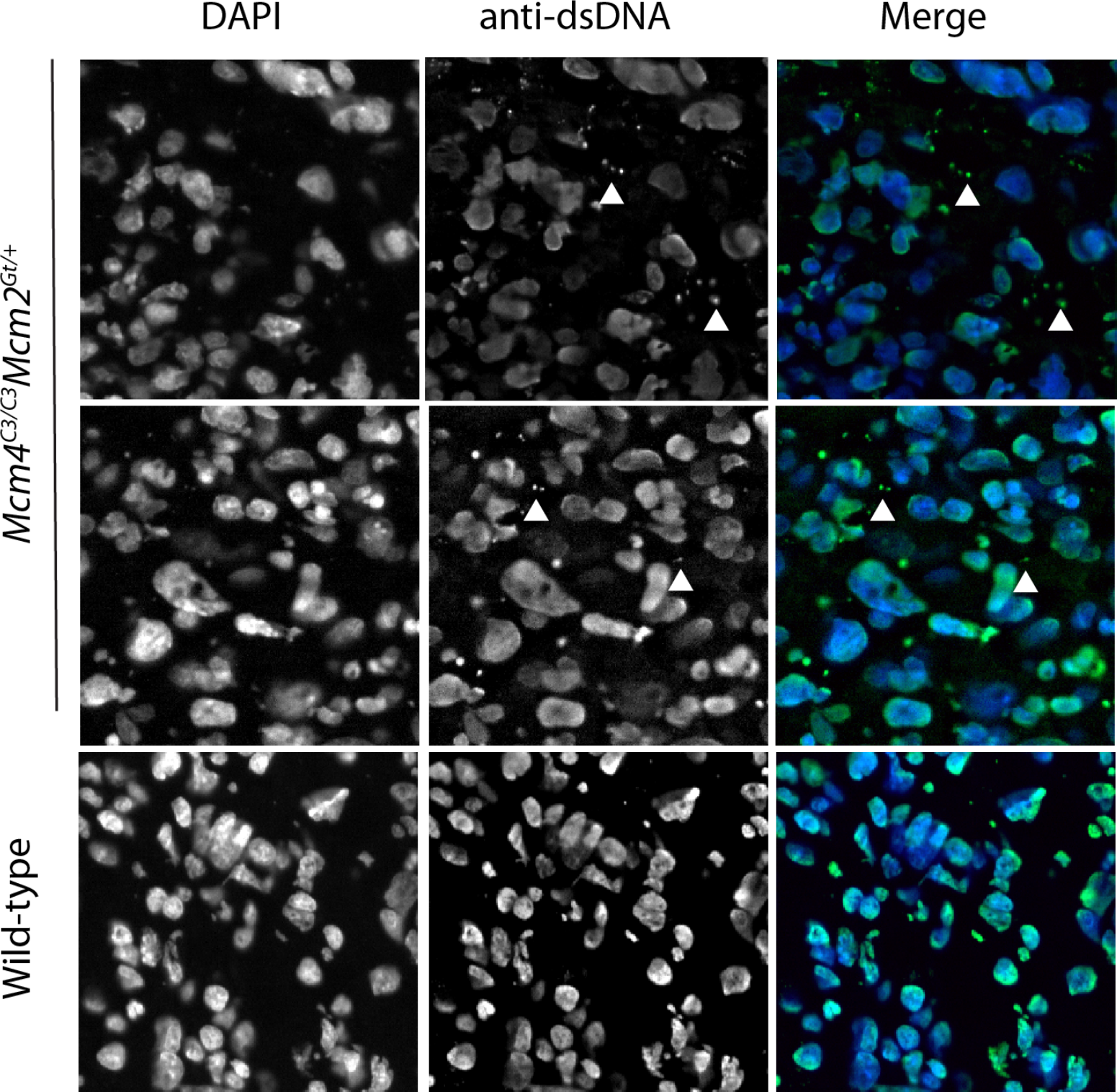
Images of E13.5 *Mcm4*^*C3/C3*^ *Mcm2*^*Gt/+*^ and wild-type placentas stained with anti-dsDNA to demonstrate the presence of extra-chromosomal DNA in the cytoplasm. Quantification is shown in Fig 4b.

**Extended Data Figure 4.**
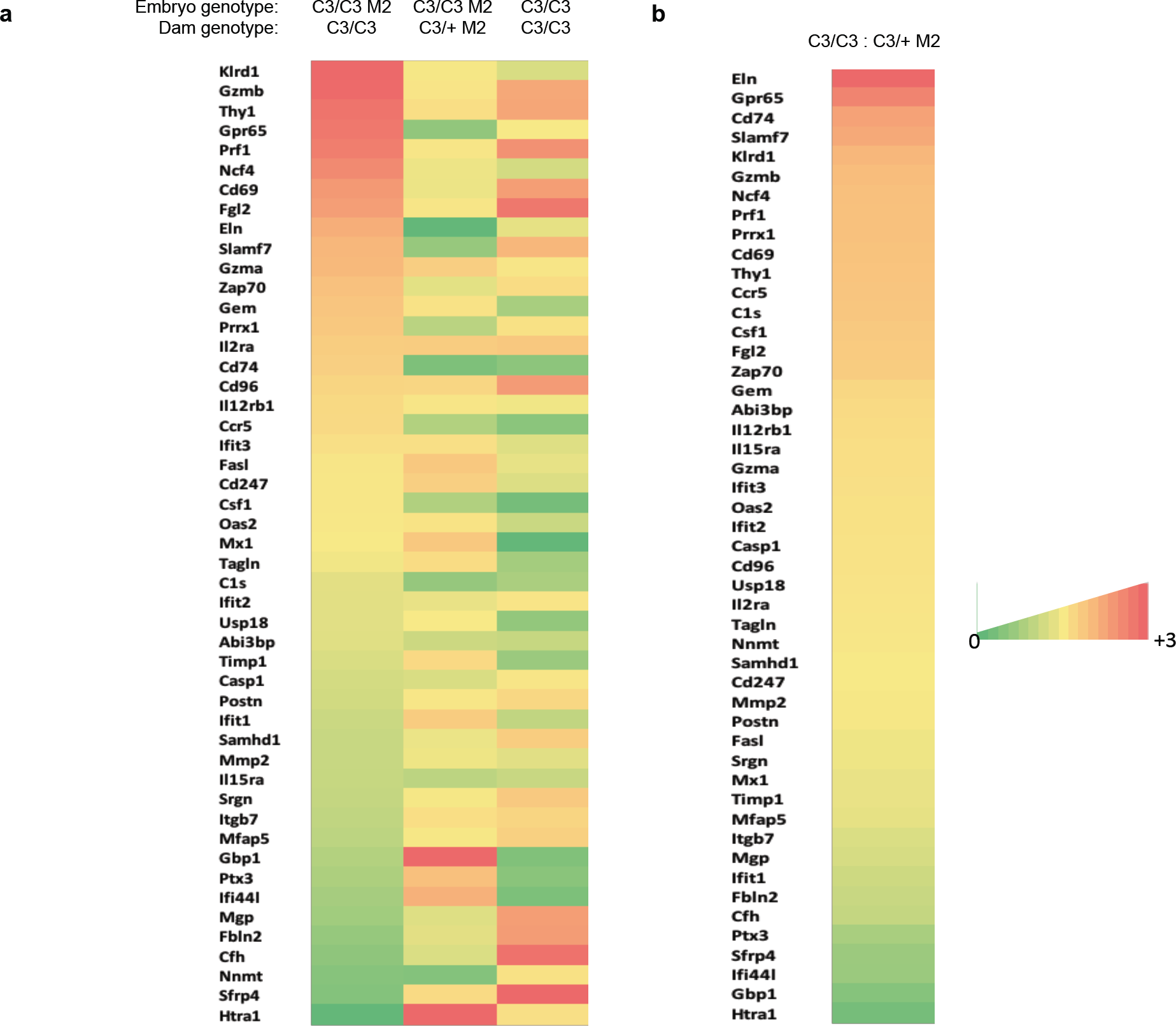
Sex specific altered expression of inflammation genes in mutants. **a**) Heatmap of the ratio of FPKM of key genes from top ranking genes from the following 3 GSEA Hallmarks: EMT, allograft rejection, and interferon gamma response. The ratios are expressed as female:male for each of the indicated embryo and dam genotypes. C3 = *Mcm4*^*C3*^; M2 = *Mcm2*^Gt/+^.**b)** Maternal genotype affects the expression of inflammation genes. Plotted are the female:male FPKM values of C3/C3 M2/+ embryos for C3/C3 dams compared to C3/+ M2 dams for the same gene sets as in (**a**). The highest and lowest genes are all related to inflammation responses.

**Extended Data Table 1.** Segregation of genotypes from crosses. a) Embryonic semilethality caused by the *Mcm4*^*C3/C3*^ *Mcm2*^*Gt/+*^ genotype is rescued by ibuprofen treatment of pregnant females. Cross: ♀ *Mcm4*^*C3/C3*^ X ♂ *Mcm4*^*C3/+*^ *Mcm2*^*Gt/+*^. Red numbers are plotted in Fig. 2A under “NSAID.” C3 = Chaos3. Data are from 29 litters. b) Embryonic semilethality caused by the *Mcm4*^*C3/C3*^ *Mcm2*^*Gt/+*^ genotype is affected by maternal genotype. Cross: ♀ *Mcm4*^*C3/+*^ *Mcm2*^*Gt/+*^ X ♂ *Mcm4*^*C3/C3*^ Red numbers are plotted in Fig. 3A under “C3/+ M2/+.” C3 = Chaos3, M2= Mcm2. Data are from 32 litters. c.) Embryonic semilethality caused by the *Mcm4*^*C3/C3*^ *Il10rb*^*-/-*^ genotype is rescued by ibuprofen treatment of pregnant females. Cross: ♀ *Mcm4*^*C3/C3*^ *Il10rb*^*+/-*^ X ♂ *Mcm4*^*C3/C3*^ *Il10rb*^*+/-*^ d.) Embryonic semi-lethality and female sex bias caused by the *FancM*^*em1Jcs/em1Jcs*^ genotype (is rescued by ibuprofen treatment of pregnant females.) Cross: ♀ *Fancm*^*em1Jcs/+*^ X ♂ *Fancm*^*em1Jcs/+*^. Red numbers are plotted in Fig. 2A.

| **a** |  |  |  |  |  |
| --- | --- | --- | --- | --- | --- |
|  | *Mcm4*<sup>C3/+</sup> | *Mcm4*<sup>C3/+</sup><br>*Mcm2*<sup>Gt/+</sup> | *Mcm4*<sup>C3/C3</sup> | *Mcm4*<sup>C3/C3</sup><br>*Mcm2*<sup>Gt/+</sup> | Total |
| **Males** | 29 | 38 | 21 | 14 | 102 |
| **Females** | 17 | 38 | 26 | 14 | 95 |
| **Total** | 46 | 76 | 47 | 28 | 197 |
| **%Female** | 36.9 | 50 | 55.3 | 50 | 48.2 |

| **b** |  |  |  |  |  |
| --- | --- | --- | --- | --- | --- |
|  | *Mcm4*<sup>C3/+</sup> | *Mcm4*<sup>C3/+</sup><br>*Mcm2*<sup>Gt/+</sup> | *Mcm4*<sup>C3/C3</sup> | *Mcm4*<sup>C3/C3</sup><br>*Mcm2*<sup>Gt/+</sup> | Total |
| **Males** | 31 | 39 | 29 | 10 | 109 |
| **Females** | 25 | 38 | 17 | 14 | 94 |
| **Total** | 56 | 77 | 46 | 24 | 203 |
| **%Female** | 44.6 | 49.4 | 36.9 | 58.3 | 46.3 |

| **c** |  |  |  |  |
| --- | --- | --- | --- | --- |
|  | *Mcm4*<sup>C3/C3</sup><br>*Il10rb*<sup>-/-</sup> | *Mcm4*<sup>C3/C3</sup><br>*Il10rb*<sup>+/-</sup> | *Mcm4*<sup>C3/C3</sup><br>*Il10rb*<sup>+/+</sup> | Total |
| **Males** | 1 | 15 | 3 | 19 |
| **Females** | 0 | 17 | 9 | 26 |
| **Total** | 1 | 32 | 12 | 45 |
| **%Female** | 0 | 53.5 | 75 | 58 |
| **+NSAID** |  |  |  |  |
| **Males** | 7 | 16 | 5 | 28 |
| **Females** | 9 | 21 | 10 | 40 |
| **Total** | 16 | 37 | 15 | 68 |
| **%Female** | 56 | 57 | 67 | 59 |

**Extended Data Table 1. Segregation of genotypes from crosses.**
| **d** |  |  |  |  |
| --- | --- | --- | --- | --- |
|  | *Fancm*<sup>em1Jcs/em1Jcs</sup> | *Fancm*<sup>em1Jcs/+</sup> | *Fancm*<sup>+/+</sup> | Total |
| **Males** | 32 | 70 | 34 | 136 |
| **Females** | 19 | 63 | 32 | 114 |
| **Total** | 51 | 133 | 66 | 250 |
| **%Female** | 37 | 47 | 48 | 46 |
| **+NSAID** |  |  |  |  |
| **Males** | 27 | 49 | 29 | 105 |
| **Females** | 24 | 43 | 27 | 94 |
| **Total** | 51 | 92 | 56 | 199 |
| **%Female** | 47 | 47 | 48 | 47 |

